# Playing with the ploidy level enables to switch on and off the strict recombination control even in the vicinity of *Brassica* centromeres

**DOI:** 10.1101/2024.02.19.580878

**Authors:** Franz Boideau, Virginie Huteau, Anael Brunet, Loeiz Maillet, Olivier Coriton, Gwenn Trotoux, Maryse Lodé-Taburel, Gwenaelle Deniot, Frédérique Eber, Marie Gilet, Julien Boutte, Jérôme Morice, Cyril Falentin, Olivier Martin, Matthieu Falque, Anne-Marie Chèvre, Mathieu Rousseau-Gueutin

## Abstract

Meiotic recombination is a key biological process in plant evolution and breeding, as it generates novel genetic diversity at each generation. However, due to its importance in chromosome segregation and genomic stability, crossovers are highly regulated in both frequency and distribution. We previously demonstrated that this strict regulation is not a fatality and that it can be naturally modified (3.6-fold increased frequency and altered distribution) in an allotriploid *Brassica* hybrid (2*n*=3*x*=29; AAC), resulting from a cross between *B. napus* (2*n*=4*x*=38; AACC) and *B. rapa* (2*n*=2*x*=20; AA). Taking advantage of the recently updated *Brassica napus* genome assembly, which now includes the pericentromeric regions, we unambiguously demonstrated that crossovers occur in these normally cold regions in allotriploids, with the presence of crossovers as close as 375 kb from the centromere. We deciphered that this modified recombination landscape (both frequency and distribution) can be maintained in successive generations of allotriploidy, with even a slight increase of crossover frequency. We also showed that this deregulated meiotic behavior may revert back to a strictly regulated one when recovering an allotetraploid progeny in the second generation. Overall, we provide here for the first time a practical and natural way to switch on and off the tight recombination control in a polyploid crop. We also discuss the potential role of this modified regulation of recombination in polyploid speciation success.

## Introduction

Meiotic recombination is a key biological process in sexually reproducing eukaryotes, as it both ensures the faithful segregation of chromosomes and generates a novel genetic diversity at each generation. It is initiated in prophase I by the formation of Double-Strand Breaks (DSBs). From about 250 DSBs occurring at each meiosis in Arabidopsis, only a few (~ 10) will form crossovers (COs), the other giving rise to non-crossovers (Mercier et al. 2015). These COs belong to two classes (I and II) depending on the meiotic proteins involved. Most formed COs (70-85%) derive from the Class I, which is controlled by proteins of the ZMM group (ZYP1, ZIP2, ZIP3, ZIP4, MSH4, MSH5 and MER3, MutLγ complex with MLH1/MLH3 and HEI10). These COs are subject to interference, a process that prevents the formation of two close-by class I COs. The few remaining COs (15-30%) derive from the Class II, which depends mainly on the MUS81 pathway and are interference insensitive (Mercier et al. 2015).

In most cases, meiotic recombination is strictly regulated, with one obligate CO per pair of homologous chromosomes to ensure the proper segregation of chromosomes, but rarely more than three (Mercier et al. 2015). Additionally, the distribution of COs is not homogeneous along the chromosomes and presents a species-specific pattern: one can have a “U” distribution with its minimum at the centromere as in *Arabidopsis thaliana*, or a rather linear increase of CO rate from the centromere to the telomere as in several crop species such as wheat, maize, barley, or oilseed rape (see Wang and Copenhaver 2018 for review). In most organisms, it has been observed that 80% of the COs were concentrated in about 25% of the genome (Choi et al. 2013, Darrier et al. 2017). This uneven distribution of COs is associated with the global chromatin organization on the chromosome. Indeed, COs mainly occur in open chromatin, which harbors a low nucleosome density, a low DNA methylation and is enriched in H3K4me3 (Choi et al. 2013, 2015, Marand et al., 2017, Lian et al. 2022). In contrast, COs are largely suppressed in the regions close to the centromeres, corresponding to the chromosomal region where the kinetochore platform assembles to allow the attachment of the spindle and the segregation of chromosomes at the opposite pole of meiocytes (McKinley and Cheeseman 2016). These centromeric regions are heterochromatic, repeat-rich, dense in nucleosomes, heavily methylated and enriched in H3K9me2 (Yelina et al. 2012, Underwood et al. 2018). Despite this tight control of meiotic recombination in most organisms, some variations may exist within a species, influencing the species evolution and adaptation (Coop and Przeworski, 2022). These differences can be the result of allele or copy number variations of certain meiotic genes, such as *HEI10* (Ziolkowski et al. 2017), but also can be related to the sex (Lenormand and Dutheil, 2005). Indeed, different CO landscapes have been observed between male and female meiosis in several species, a phenomenon referred as heterochiasmy (Sardell and Kirkpatrick, 2020, Capilla-Perez et al. 2021, Cai et al. 2023).Within the last ten years, several methods have emerged to greatly increase the number of COs per meiosis, either by over-expressing pro-COs genes such as *HEI10* (Ziolkowski et al. 2017), or by knocking-out (KO) anti-COs genes (e.g. *FANCM*, *FANCC*, *RECQ4*, *FIGL1* or *ZYP1*: Crismani et al. 2012, Séguéla-Arnaud et al. 2015, Girard et al. 2015, Durand et al. 2022, Singh et al. 2023, Capilla-Perez et al. 2024) or by combining both approaches (Serra et al. 2018). As an example, the recent overexpression of the pro-COs HEI10 protein combined with the KO of the *ZYP1* gene involved in the synaptonemal complex gave rise to a massive increase of COs, but only in regions where COs normally occur (Durand et al. 2022). To date, very few cases have led to an increase of CO numbers near the centromeric regions. Most of them were obtained when performing a K.O. of genes involved in epigenetic marks. As an example, the loss of CG methylation via the KO of *MET1* (DNA hypomethylated methyltransferase 1) or *DDM1* (decreased DNA methylation 1) in Arabidopsis altered the CO distribution, with an increased COs number around the centromeres but also a decrease in pericentromeric regions (Melamed-Bessudo and Levy, 2012, Yelina et al. 2012). Similarly, the disruption of Arabidopsis H3K9me2 and non-CG DNA methylation pathways (via the mutation of *KYP*/*SUVH5*/*SUVH6*, and *CMT3*), slightly increased meiotic recombination in proximity to the centromeres, including pericentromeres (Underwood et al., 2018). A third factor affecting recombination corresponds to the ploidy level. In the case of allopolyploid species, corresponding to the presence of at least two different genomes in the same nucleus, it has been observed in some species a 2-fold increase of recombination rate, as exemplified in *Gossypium* (Brubaker et al. 1999, Desai et al. 2006), Arabidopsis (Pecinka et al. 2011), Brassicoraphanus (Park et al. 2020), or wheat (Wan et al., 2021, Yang et al. 2022). In *Brassica*, it has been found that allotriploidy increased by 3.5-fold the homologous recombination compared to its diploid and even its allotetraploid counterpart, with also 1.7 times more recombination in female meiosis compared to the male meiosis (Pelé et al. 2017). Interestingly, allotriploids present a modified recombination pattern, independently of the sex of meiosis (Leflon et al. 2010, Pelé et al. 2017, Boideau et al. 2021). Within such allotriploid *Brassica* hybrids (2*n*=3*x*=29; AAC) deriving from a cross between *B. napus* (2*n*=4*x*=38; AACC) and *B. rapa* (2*n*=2*x*=20; AA), the A chromosomes pair as ten bivalents, whereas the remaining nine C chromosomes remain as univalent at metaphase I and segregate randomly within gametes (Leflon et al. 2006). It has been proposed that the presence of univalents may cause the nucleus to linger in a recombination active state, resulting in an increase CO number and in a modified distribution in the remaining chromosomes that are correctly synapsed (Martinez-Perez and Moore, 2008). Until recently, it remained unknown if this unique recombination was specific to *Brassica* and allotriploids. However, a similar study had been performed in a monocotyledon pentaploid wheat, deriving from a cross between a hexaploid wheat (*Triticum aestivum*, 2*n*=6*x*=42, AABBDD) and a tetraploid wheat (*Triticum turgidum*, 2*n*=4*x*=28). That study also found a CO increase (3 to 4-fold) between either A or B homologus chromosomes and a modified distribution (Yang et al. 2022). These different results suggest that this modified recombination pattern observed in both a monocotyledon and dicotyledon may be used to efficiently improve the genetic diversity and breeding of many polyploid crops, such as wheat, cotton, oilseed rape, coffee and strawberry (Leitch and Leitch, 2008).

In this study, our first aim was to investigate to what extent allotriploidy may facilitate the formation of COs near the cold recombining centromeric regions, by taking advantage of the recently updated genome assembly of the *B. napus* cv. Darmor genome (Rousseau-Gueutin et al. 2020) that now contains the repeat rich (peri)centromeric regions. Additionally, it remained to be deciphered if this modified recombination pattern may be kept or may revert back to a normal strict meiotic behavior to prevent putative long-term genomic instabilities. To address these questions, we firstly designed numerous markers specific to two pericentromeric regions. Thereafter, we took advantage of the formation of some allotriploid (AAC) and allotetraploid (AACC) individuals within the progeny of an AAC allotriploid hybrid, to investigate the impact of the ploidy level in the following generation on homologous recombination. Overall, we produced and genotyped a total of six segregating populations (different ploidy levels and different generations). To compare the recombination frequency and distribution between these hybrids, we performed comparative genetic mapping using the same anchored SNPs set (including pericentromeric markers), and always validated them by HEI10 immunostaining on Pollen Mother Cells (PMC) associated with GISH-like. From these analyses, we demonstrated that (1) COs may occur as close as 375 kb from the centromere in an allotriploid compared to 4.5 Mb in an allotetraploid, (2) the modified recombination landscape can be maintained in successive generations of allotriploidy, (3) a normal meiotic control is recovered in an allotetraploid progeny of an allotriploid hybrid, (4) the recombination rate is increased within hybrids of the second generation, whatever their ploidy levels. This novel knowledge is important for both fundamental and applied research, as it increases our understanding on how allotriploidy may play a major role in polyploid speciation, evolution and adaptation. In addition, the method described here provides an efficient and easy way to naturally switch on and off a strict control of recombination in polyploid crops, that can be used to greatly enhance their genetic diversity, reduce linkage disequilibrium for some agronomic traits and favor the combination of favorable alleles, or even facilitate the identification of candidate genes of interest.

## Results

### 1. Meiotic behaviors, pollen viability and seed set within hybrids presenting varying ploidy levels

As an initial step to analyzing the evolution over successive generations of the recombination profile in *Brassica* hybrids harboring different ploidy levels, we investigated the meiotic behavior at metaphase I of an allotetraploid (A_n_A_r_C_n_C_o_) and allotriploid (A_n_A_r_C_n_) F1 hybrids presenting the exact same A genotype, as well as their backcross progenies. To facilitate the reading, we chose the following nomenclature to describe our hybrids, with first the ploidy level, then the generation, and finally the maternal ploidy origin of each hybrid. Briefly, the F1 allotetraploid and allotriploid hybrids are referred to 4x-F1 and 3x-F1, respectively. The descending allotetraploid and allotriploid hybrids deriving from the backross of the 3x-F1 were entitled 4x-B1F1^3x^ and 3x-B1F1^3x^, whereas the allotetraploid hybrid deriving from the backcross of the 4x-F1 was referred to as 4x-B1F1^4x^. More details on the genotypes and on the methods used to create these hybrids is given in Figure 1.

**Figure 1:**
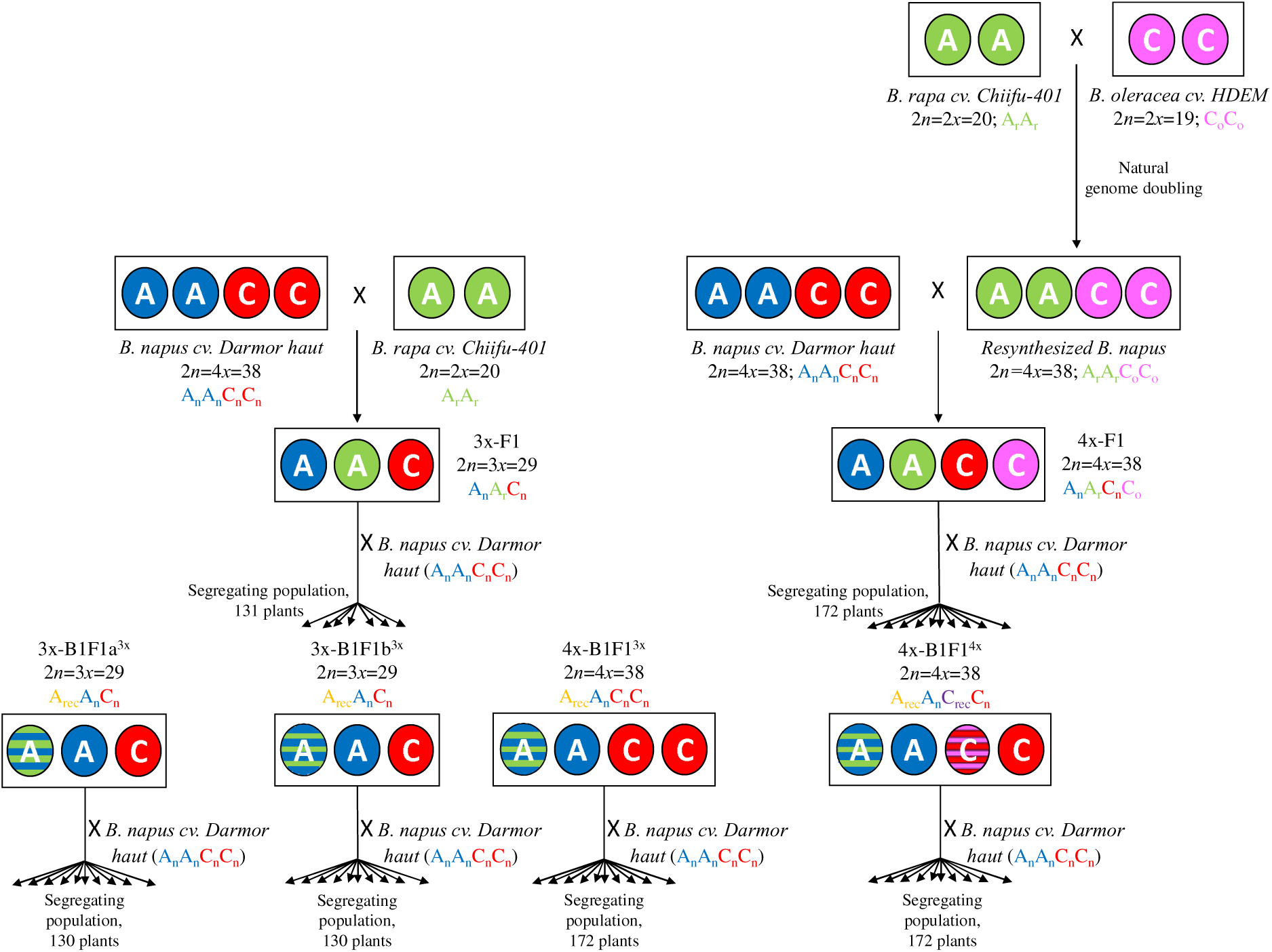
Schematic representation of the production of the F1 and B1F1 hybrids and their corresponding segregating backcross populations. The names and the ploidy level of the parental lines are indicated and corresponds to *B. napus* cv. Darmor (2*n*=4*x*=38, A_n_A_n_C_n_C_n_), to *B. rapa cv.* Chiffu (2*n*=2*x*=20; A_r_A_r_) and *B. oleracea* cv HDEM (2*n*=2*x*=18; C_o_C_o_). The nomenclature A_r_ and A_n_ correspond to the A genome of *B. rapa* and *B. napus*, respectively, whereas C_o_ and C_n_ refer to the C genome of *B. oleracea* and *B. napus*, respectively. The index “rec” indicates that recombination occurred on the genome concerned. The allotriploid lineage is present on the left part of the diagram, while the right part corresponds to the allotetraploid lineage. The number of genotyped progenies for each of the six investigated segregating populations is indicated.

For each hybrid, the meiotic behavior was firstly established using both classical aceto-carmine staining and a GISH-like technique using the Bob014O06 BAC probe that specifically hybridizes to all C chromosomes. The 3x-F1 and 4x-F1 hybrids showed the expected meiotic behavior, with a majority of Pollen Mother Cells (PMC) with ten A bivalents plus nine C univalents for 3x-F1 and ten A bivalents plus nine C bivalents 4x-F1 (Table 1, Figure 2). We also assessed the pollen viability and seed set of these hybrids and observed that the 3x-F1 presents a lower pollen fertility compared to the 4x-F1 hybrid (69% and 97%, respectively, Table 1). Similarly, there is a lower number of seeds per pollinated flower in the 3x-F1 compared to the 4x-F1 (2.3 vs 10.2, respectively, Table 1). At the second generation, we chose to study two different 3x-B1F1^3x^ hybrids carrying different heterozygous genomic regions in order to validate that whatever the genetic structure of the hybrid we would observe the same meiotic recombination pattern. The two selected 3x-B1F1^3x^ hybrids had an improved meiotic behavior compared to the 3x-F1, with systematically ten A bivalents and nine C univalent. They presented a male fertility and seed set similar to their parental mother plant 3x-F1 (Table 1, Figure 2). Concerning the 4x-B1F1^3x^ and 4x-B1F1^4x^, they also both presented a similar meiotic behavior, pollen viability and seed set as their mother 4x-F1 plant (Table 1, Figure 2).

**Figure 2:**
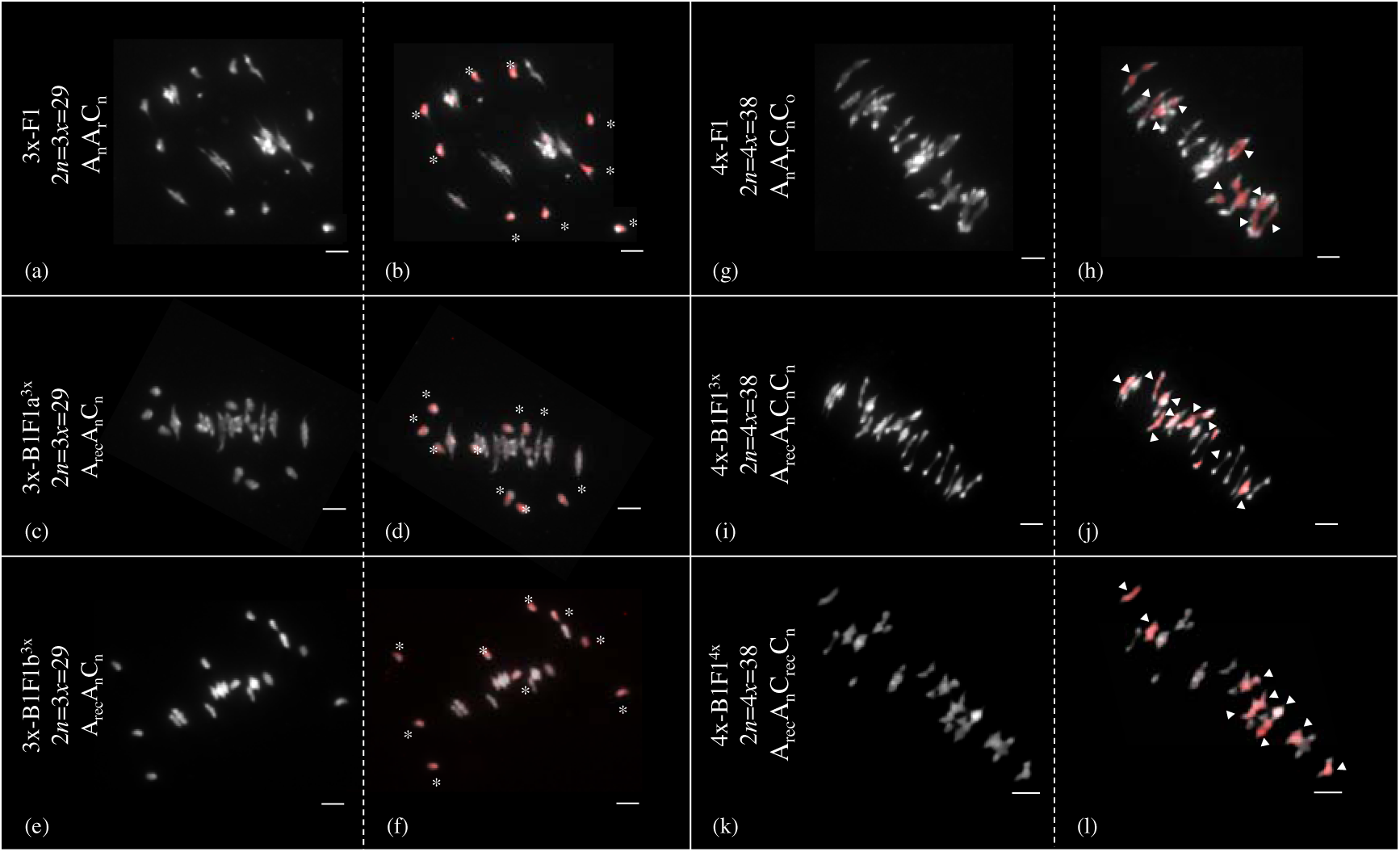
Meiotic behavior on chromosome spreads in metaphase I in the different allotriploid and allotetraploid hybrids using the C genome specific BoB014O06 probe. Chromosomes in grey are from the A-genome and chromosomes painted in red are from the C-genome. GISH-like analyses of meiotic chromosomes were carried out in the allotriploid (left) and allotetraploid (right) hybrids: 3x-F1 (A_n_A_r_C_n_) (a-b), 3x-B1F1a^3x^ (A_rec_A_n_C_n_) (c-d), 3x-B1F1a^3x^ (A_rec_A_n_C_n_) (e-f), 4x-F1 (A_n_A_r_C_n_C_o_) (g-h), 4x-B1F1^3x^ (A_rec_A_n_C_n_C_n_) (i-j), 4x-B1F1^4x^ (A_rec_A_n_C_rec_C_n_) (k-l). The nine C univalent in allotriploids (b, d, f) are indicated by stars, whereas the nine C bivalents in allotetraploids are shown with white arrows (h, j, l). Bars = 5µm. Original pictures from the 3x-F1 and 4x-F1 (a, b, g, h) derived from Boideau et al. 2021.

**Table 1:** Meiotic behavior, pollen fertility and seed set of the hybrids used in this study.

| Hybrid | Genomic structure | Number of PMC | Meiotic behavior | Percentage of cells at the expected meiotic behavior | Pollen viability (%) | Average seed set per pollinated flower | References |
| --- | --- | --- | --- | --- | --- | --- | --- |
| 3x-F1 | $A_n A_r C_n$,<br>$2n=3x=29$ | 21 | 9,38I+9,81II | 81 | 68.80 | 2.26 | Boideau et al. 2020 |
| 4x-F1 | $A_n A_r C_n C_o$,<br>$2n=4x=38$ | 20 | 0,2I+18,8II+0,05IV | 85 | 96.70 | 10.16 | Boideau et al. 2020 |
| 3x-B1F1a<sup>3x</sup> | $A_{rec} A_n C_n$,<br>$2n=3x=29$ | 20 | 9I +10II | 100 | 64.90 | 1.52 | This study |
| 3x-B1F1b<sup>3x</sup> | $A_{rec} A_n C_n$,<br>$2n=3x=29$ | 20 | 9I+10II | 100 | 61.40 | 1.49 | This study |
| 4x-B1F1<sup>3x</sup> | $A_{rec} A_n C_n C_n$,<br>$2n=4x=38$ | 17 | 0,2I+18,8II+0,05IV | 85 | 93.80 | 13.22 | This study |
| 4x-B1F1<sup>4x</sup> | $A_{rec} A_n C_{rec} C_n$,<br>$2n=4x=38$ | 20 | 0.3I + 18.75II + 0.05IV | 80 | 96.70 | 14.42 | This study |

In the following sections, we compared the recombination profile of these different hybrids by performing genetic mapping using an identical set of polymorphic SNP markers for each comparison. These results were thereafter complemented by the quantification of class I COs frequency on the A sub-genome in PMC (male meiosis) at diakinesis, using HEI10 immunostaining coupled with GISH-like chromosome painting using the C-chromosome specific BAC.

### 2. Allotriploidy increases recombination frequency and enables the formation of COs at the vicinity of centromeres, notably via deeply reducing the strength of interference

Backcross populations of the 4x-F1 and 3x-F1 hybrids (always used as females) were previously obtained and genotyped using *Brassica* Illumina Infinium SNP array (SGS-TraitGenetics GmbH, Gatersleben, Germany). However, the recently improved *B. napus cv. Darmor* genome using the third-generation sequencing technologies (Rousseau-Gueutin et al. 2020) enabled to assemble the complex and repetitive-rich pericentromeric regions (Figure S1) and revealed that almost no markers were in fact present in these regions. Indeed, in the previous study, the closest markers surrounding the A01 and A02 centromeres were previously distant by 5 Mb and 7.6 Mb and thus did not allow to identify the presence of COs in these regions. To circumvent this problem and properly investigate the presence of COs near the centromere, we took advantage of the recent improvements of both *B. napus* cv. Darmor and *B. rapa* cv. Chiifu genome assemblies (A parental donors of our hybrids) to design many novel polymorphic KASPar markers in both A01 (81 markers) and A02 (25 markers) pericentromeric regions (Table S1). These two chromosomes were chosen as that were polymorphic in most of the hybrids investigated in this study, including most second-generation hybrids (3x-B1F1b^3x^, 4x-B1F1^3x^, 4x-B1F1^4x^, see Figure 1).

Using the same set of polymorphic markers deriving from both the 15K array and these newly designed KASPar markers (totalling 2274 SNPs), we generated genetic maps for both 3x-F1 and 4x-F1. As previously observed (Boideau et al. 2021), the 3x-F1 hybrid shows a significant 3.7-fold increase of COs frequency compared to the 4x-F1 hybrid (Bonferroni corrected chi-squared test (BcC test), P < 2.2E-16, Table S2).

When investigating more finely the pericentromeric regions, we found that the 4x-F1 hybrid had no CO, whereas the 3x-F1 hybrid presented 36 COs in total (Table S2). Interestingly, these COs can arise as close as 375 kb from the centromeric borders in the 3x-F1, while the nearest CO detected for the 4x-F1 hybrid was localized 4.5 Mb from the centromeric border. We also compared the strength of interference between these hybrids and observed that the level of interference is significantly lower in the 3x-F1 compared to the 4x-F1 hybrid, with an estimated Nu value of 1.847 and 9.588 respectively. Nevertheless, a slight interference was still detected in the 3x-F1 hybrid according to the gamma model (testing if Nu was close to 1, P < 0.001).

The increased CO frequency in the 3x-F1 compared to 4x-F1 hybrids observed in female meiosis using genetic mapping was thereafter validated using HEI10 immunostaining, labelling class I COs on male meiocytes at diakinesis (24 in the 3x-F1 vs 18 in the 4x-F1 on average, Mann Whitney Wilcoxon test (MWW test), P = 2.05E-08, Figures 3 and 4, Table S3).

**Figure 3:**
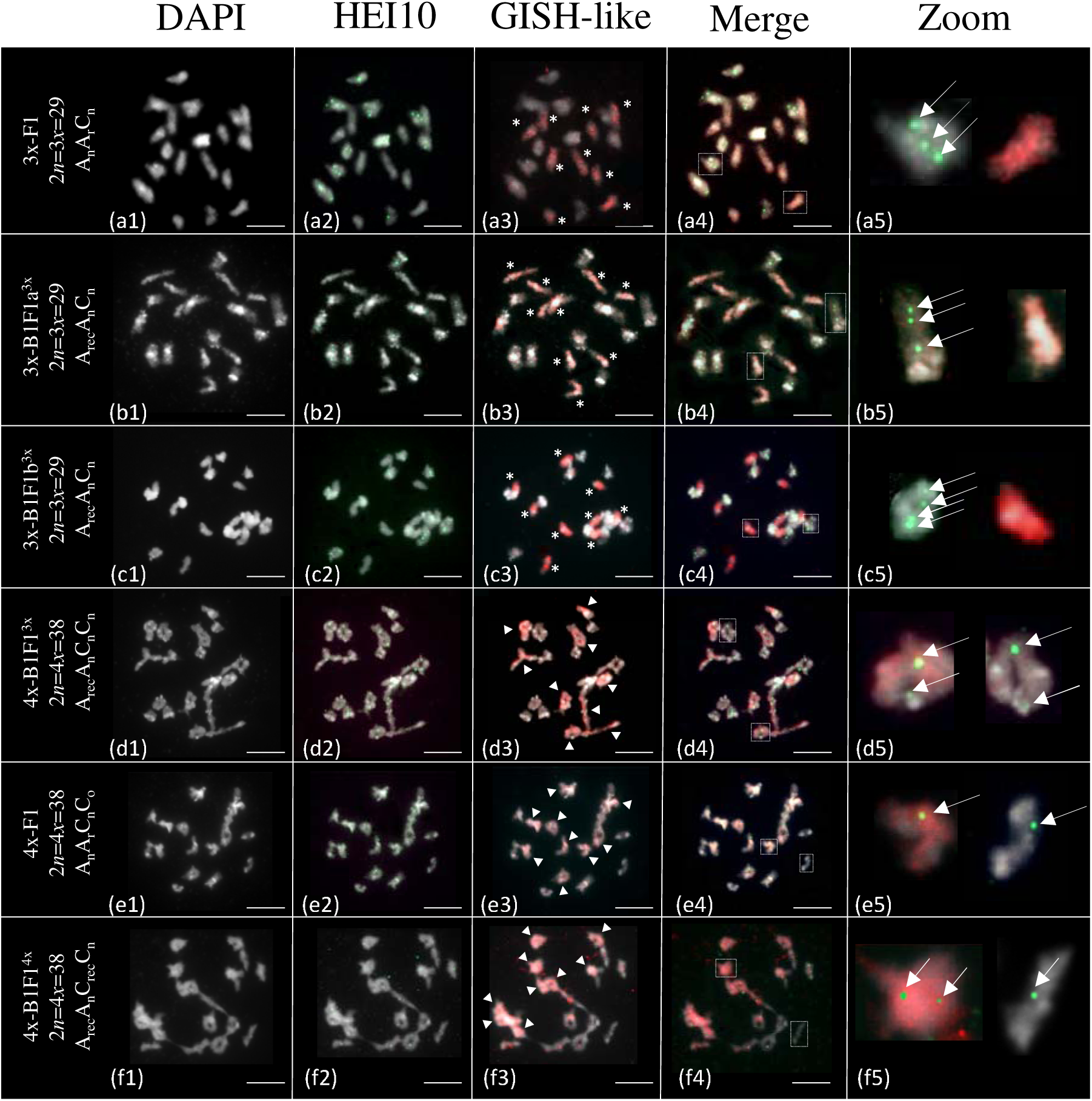
Immunolocalization of HEI10 and GISH-like labelling on diakinesis chromosomes in pollen mother cells of the various allotriploid (a, b, c) and allotetraploid (d, e, f) hybrids. The investigated genotypes are: 3x-F1 (A_n_A_r_C_n_) (a), 3x-B1F1a^3x^ (A_rec_A_n_C_n_) (b), 3x-B1F1b^3x^ (A_rec_A_n_C_n_) (c), 4x-B1F1^3x^ (A_rec_A_n_C_n_C_n_) (d), 4x-F1 (A_n_A_r_C_n_C_o_) (e) and 4x-B1F1^4x^ (A_rec_A_n_C_rec_C_n_) (f). For each cell, chromosomes were counterstained with DAPI (white, a1-f1), HEI10 immunolabeled (green, a2-f2) and red painted chromosomes derive from the C-subgenome (BoB014O06, a3-f3). The overlay of the three signals is shown in the column “Merge” (a4-f4) and a focus on some A and C chromosomes highlighted with a dotted white rectangle is displayed at the end of each rows in the column “Zoom” (a5-f5). In the third column, the nine C univalent chromosomes in the allotriploids are indicated by stars (a3, b3, c3) and the ten C bivalents in the allotetraploid hybrids are indicated by white arrow heads (d3, e3, f3). HEI10 foci are highlighted by white arrows in the last column (a5-f5). Bars = 5µm.

**Figure 4:**
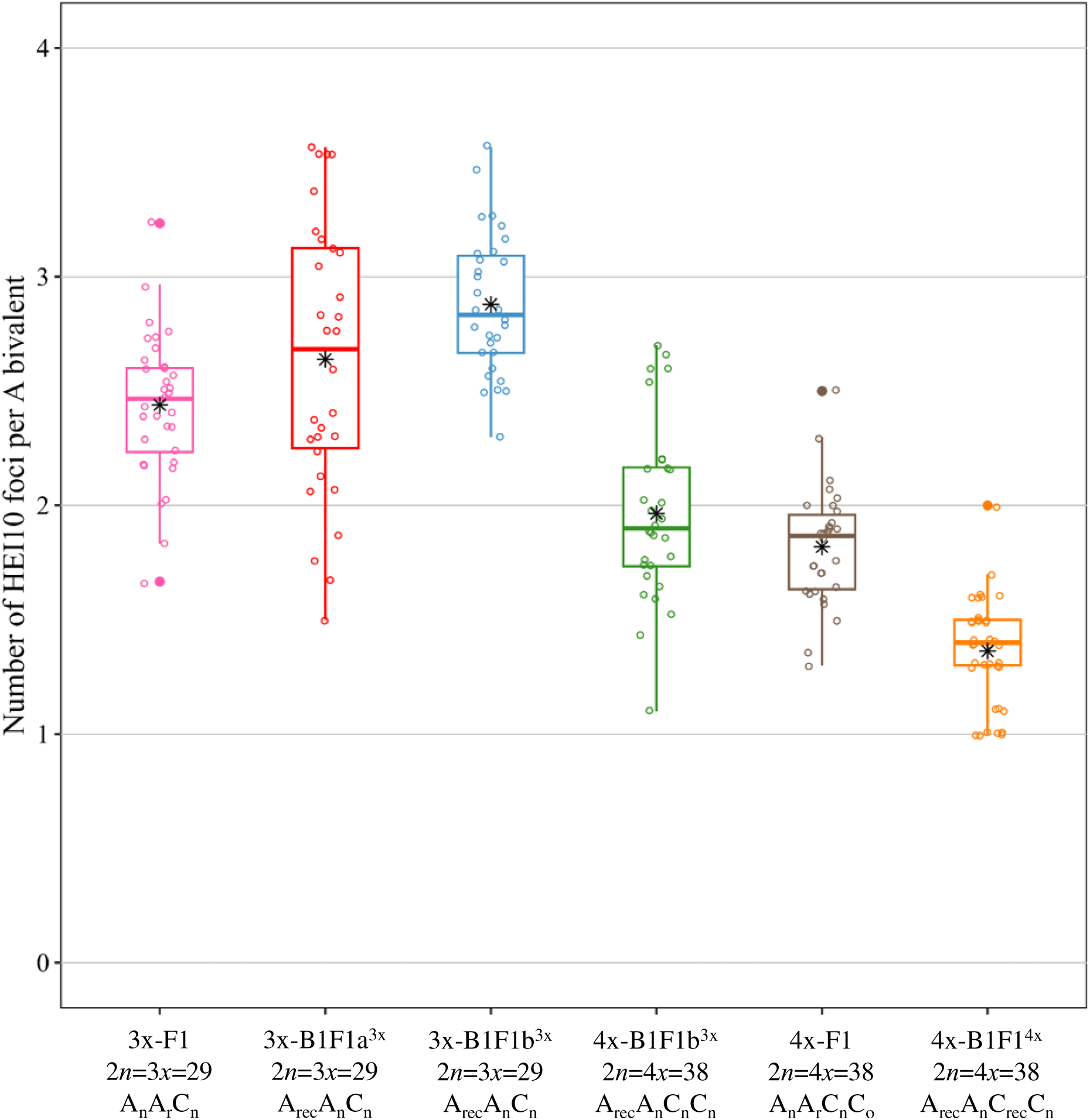
Average number of HEI10 foci per A bivalent in the hybrids. The six investigated hybrids are as following: 3x-F1 (A_n_A_r_C_n_), 3x-B1F1a^3x^ (A_rec_A_n_C_n_), 3x-B1F1b^3x^ (A_rec_A_n_C_n_), 4x-B1F1^3x^ (A_rec_A_n_C_n_C_n_), 4x-F1 (A_n_A_r_C_n_C_o_) and 4x-B1F1^4x^ (A_rec_A_n_C_rec_C_n_). The means are indicated by a black asterisk.

### 3. The modified recombination profile observed in AAC allotriploids can be maintained by performing successive generations at the allotriploid level with even a slight increase of CO frequency

To determine if the modified recombination landscape observed in the 3x-F1 may be conserved for successive generations at the allotriploid level, we searched for allotriploid plants in the backcross progeny of the 3x-F1 using firstly flow cytometry. We identified 8 out of the 245 genotyped progenies that were potentially allotriploids. Their chromosome number were thereafter validated at 2*n*=3*x*=29 using molecular cytogenetics. It is important to note that as these progenies were obtained via a backcross, on average only half of the A genome is expected to be heterozygous. For the following analyses, we thus kept one 3x-B1F1b^3x^ hybrid, which was heterozygote for the A01 and A02 pericentromere regions, allowing a comparison of these regions in the successive generations. Comparative genetic mapping was only performed in the common heterozygous regions between the 3x-B1F1b^3x^ and 3x-F1 hybrids (distributed in 18 regions of the ten A chromosomes, totaling 182.3 over the 346 Mb) using the same set of 1205 polymorphic SNP markers (Table 2, Figure 5), thus preventing biases when comparing their recombination profiles. It revealed that the 3x-B1F1b^3x^ presents a similar recombination profile with even a statistically significant 1.7-fold increase of recombination frequency compared to the 3x-F1 hybrid (BcC test, 3.82E-78). This significant increase was observed for non-pericentromeric and pericentromeric regions (BcC test, P = 1.36E-14, 2.71E-65 respectively). It was also validated in male meiosis using HEI10 immunostaining (1.18-fold increase, MWW test, P=1.27E-05, Figure 3 and 4).

**Figure 5:**
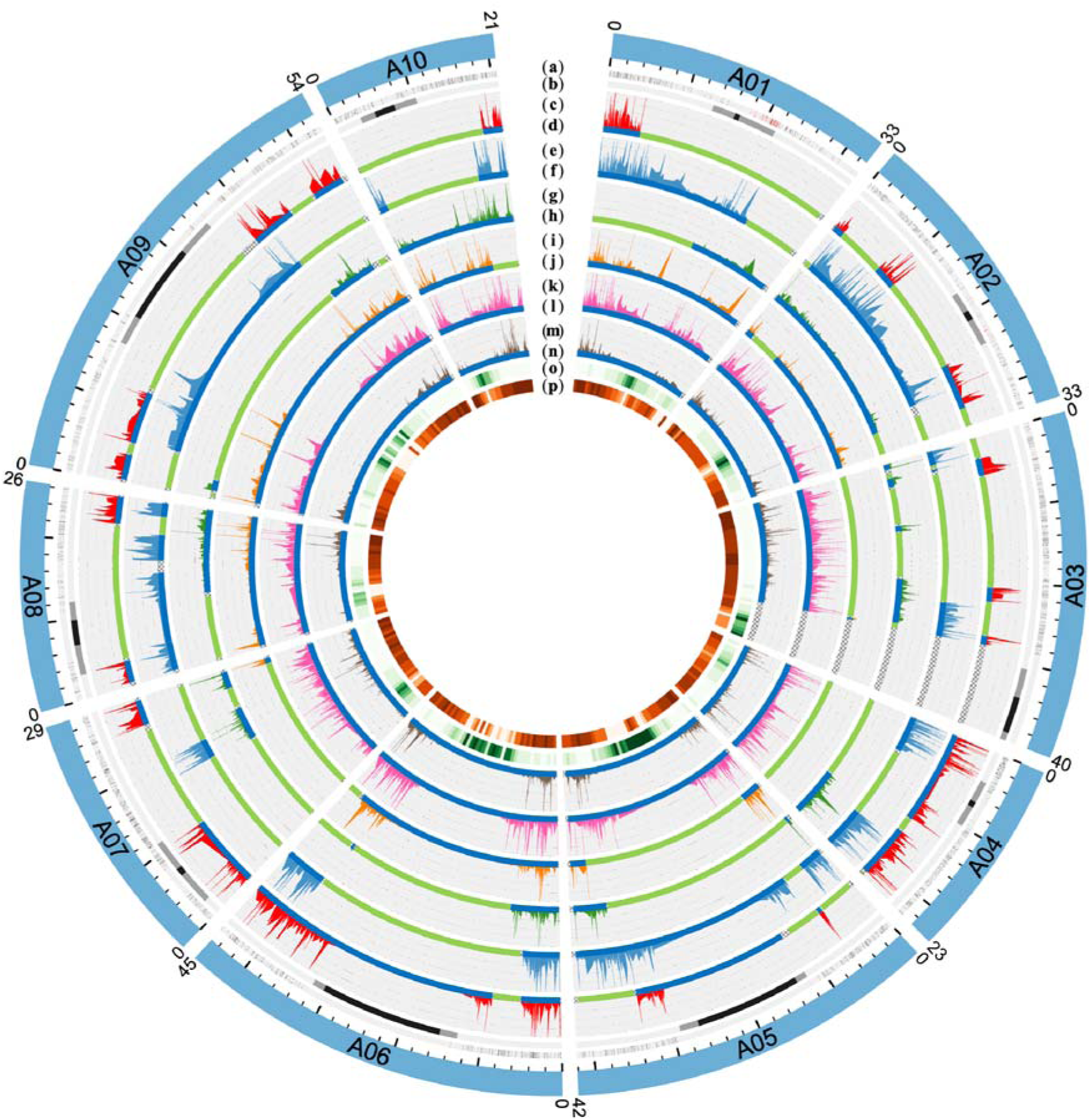
Homologous recombination landscape on the A genome of the six hybrids. The first outer circle represents the ten A chromosomes from *B. napus* cv. Darmor-bzh. The second circle represents the position of the polymorphic SNPs used to generate the genetic maps (a). The red polymorphic SNPs on chromosomes A01 and A02 correspond to the pericentromeric KASPar developed in this study. The third circle refers to the position of the pericentromeric and centromeric regions in grey and black respectively (b). The lineplots correspond to the recombination rates in the 3x-B1F1a^3x^ (c), 3x-B1F1b^3x^ (e), 4x-B1F1^3x^ (g), 4x-B1F1^4x^ (i), 3x-F1 (k) and 4x-F1 (m) hybrids. The genotyping blocks bellow each lineplot indicate that the region is either heterozygous (blue) or homozygous (green) in the 3x-B1F1a^3x^ (d), 3x-B1F1b^3x^ (f), 4x-B1F1^3x^ (h), 4x-B1F1^4x^ (j), 3x-F1 (l) and 4x-F1 (n) hybrids. The hatched areas (as example: end of chromosome A03) correspond to regions without informative genotyping data. Finally, the two most inner circles represent the transposable elements density (o) and gene density (p), respectively.

**Table 2:** Characterization of the hybrids and segregating populations. For each hybrid and associated segregating population, several parameters are indicated. The star indicate that the F1 hybrid is fully heterozygous for the A subgenome

| Hybrid | Genomic structure | Number of plants | Number of heterozygous regions | Number of polymorphic markers | Number of COs | Genetic size (cM) | Genome coverage (Mb) | Genome coverage (%) | Average recombination rate (cM/Mb) |
| --- | --- | --- | --- | --- | --- | --- | --- | --- | --- |
| 3x-F1 | $A_n A_r C_n$,<br>$2n=3x=29$ | 131 | 10* | 2274 | 3946 | 3045.3 | 327.58 | 94.55 | 9.30 |
| 3x-B1F1a<sup>3x</sup> | $A_{rec} A_n C_n$,<br>$2n=3x=29$ | 130 | 22 | 980 | 2216 | 1739.7 | 146.68 | 42.33 | 11.86 |
| 3x-B1F1b<sup>3x</sup> | $A_{rec} A_n C_n$,<br>$2n=3x=29$ | 130 | 18 | 1205 | 3365 | 2673.3 | 182.27 | 52.61 | 14.67 |
| 4x-B1F1<sup>3x</sup> | $A_{rec} A_n C_n C_n$,<br>$2n=4x=38$ | 172 | 17 | 1128 | 946 | 574.6 | 125.18 | 36.13 | 4.59 |
| 4x-F1 | $A_n A_r C_n C_o$,<br>$2n=4x=38$ | 172 | 10* | 2274 | 1408 | 829.6 | 327.58 | 94.55 | 2.53 |
| 4x-B1F1<sup>4x</sup> | $A_{rec} A_n C_{rec} C_n$,<br>$2n=4x=38$ | 172 | 14 | 1099 | 1093 | 665.4 | 198.84 | 57.39 | 3.35 |

To further validate this increased recombination frequency in the 3x-B1F1b^3x^ compared to the 3x-F1, similar experiments were performed on another 3x-B1F1a^3x^ plant. Comparative genetic maps also showed a significant increase of recombination frequency (1.35 folds, BcC test, P=1.92E-19, Table 1, Figure 5, Table S2) compared to the 3x-F1, while HEI10 immunostaining revealed a slight but not significant increase of HEI10 foci (1.08-fold, MWW test, P=0.98, Figure 3 and 4, Table S3). Therefore, the modified recombination profile observed in the 3x-F1 is maintained in the 3x-B1F1 hybrid, with even a slight increase of the recombination frequency.

### 4. The deregulation of the recombination control provoked by allotriploidy can be switched off and reverted back to normal in a descending allotetraploid individual

To decipher if the modified recombination landscape observed in the 3x-F1 hybrid may revert back to a normal and strict meiotic control in the next generation, we firstly searched for an allotetraploid individual in the backcross progeny of the 3x-F1 using both flow cytometry and molecular cytogenetics. In total, we identified two potential allotetraploid progenies and we chose one individual (referred as 4x-B1F1^3x^, Figure 1) that was heterozyogous for the A01 and A02 pericentromeric regions. In this 4x-B1F1^3x^ plant, we could identify 17 polymorphic regions, totaling as expected about half of the A genome. We performed comparative genetic mapping between the 4x-B1F1^3x^ and 3x-F1 (Figure 5, Table 2) and observed a significant 2.6-fold decrease of the recombination frequency in the 4x-B1F1^3x^ compared to the 3x-F1 hybrid (BcC test, P = 2.45E-19, Table 1, Table S2). A significant decrease of class I COs was also observed in male meiosis using HEI10 immunostaining (1.24-fold, MWW test, P = 5.92E-05, Figure 3 and 4, Table S3).

Interestingly, using both comparative genetic mapping or HEI10 immunostaining, the decrease of the recombination frequency observed in the 4x-B1F1^3x^ was smaller than expected, compared to the observed 3.7-fold observed between the 4x-F1 and the 3x-F1. The smaller decrease of the recombination frequency in the 4x-B1F1^3x^ can either be due to a residual effect from the allotriploid step or to its backcross origin, with only half of the genome that contains heterozygous regions juxtaposed to homozygous regions. To test these hypotheses, we compared the recombination frequency between a 4x-B1F1^4x^ and a 4x-F1 hybrid using both genetic mapping and HEI10 immunostaining (Table 2, Figure 4). We observed an increased recombination frequency in the 4x-B1F1^4x^ compared to the 4x-F1, as previously observed for the allotriploid lineage. We also compared the recombination frequency between a 4x-B1F1^3x^ and 4x-B1F1^4x^ using the same set of markers, and surprisingly observed slightly more recombination in the 4x-B1F1^4x^ compared to the 4x-B1F1^3x^ (BcC test, P = 0.026). However, this difference was only due to one region on the chromosome A07 (BcC test, P = 0.085). Altogether, these results highlight an absence of residual effect from the allotriploid step and suggests that the increased recombination frequency observed in the second generation is a result of the backcrossing.

## Discussion

In this study, we further investigated the modified recombination pattern discovered in *Brassica* allotriploids. First, we determined unambiguously that the intriguing recombination profile observed in allotriploids enables the formation of numerous COs in the normally low recombining pericentromeric regions. Secondly, we identified individuals of different ploidy levels (either allotriploid or allotetraploid) at the following generation and determined that it was either possible to naturally maintain this deregulation of recombination or to revert it back to a normal strict meiotic control. Finally, we discuss the potential origins of this meiotic deregulation and the important role that allotriploidy may play in polyploid diversification and establishment.

### Allotriploidy massively increases the CO formation in the normally low recombining pericentromeric regions

In spite of the hypothesis that the absence of COs in centromeric regions may be related to the key role of this chromosomic region in the proper segregation of chromosomes (McKinley and Cheeseman, 2016), we demonstrated that they occur in natural allotriploid hybrids at a high frequency in pericentromeric regions. Due to their key cellular function, the centromeric regions, which are composed of megabase of satellite repeat arrays (Naish et al. 2021), are lacking COs (Fernandes et al. 2024). Even large regions (several Mb) surrounding the centromeres present a very low recombination rate, as recently exemplified in Arabidopsis (Fernandes et al. 2024) or also in crops, such as barley (Dreissig et al. 2020) and wheat (Raz et al. 2021). Here, we took advantage of the recent improvements of *B. napus* cv. Darmor (Rousseau-Gueutin et al. 2020) and *B. rapa* cv. Chiifu (Zhang et al. 2018) genome assemblies, which now includes the pericentromeres, to determine if allotriploidy facilitates the formation of COs in these latter regions. Indeed, our previous study (Pele et al. 2017) that used the first *B. rapa* genome assembly (Wang et al. 2011) did in fact not contain any pericentromeric marker from these regions, preventing us to truly test this hypothesis. To that purpose, we designed numerous novel KASPar markers specific to the pericentromeric regions of chromosomes A01 and A02 spanning 8.2 Mb and 7.1 Mb, respectively, and that were heterozygous in most of our investigated hybrids (five out of six). This marker enrichment enabled us to demonstrate that numerous COs occur in these normally cold pericentromeres in allotriploids as 36 out of 131 individuals had a CO compared to none in an allotetraploid population of 172 individuals. Additionally, COs may occur as close to 375 kb from the centromere border in allotriploids compared to 4.5 Mb in allotetraploids. In plants, the quasi absence of COs in the pericentromeric regions correlate with the presence of high DNA methylation and an enrichment for heterochromatic histone marks, such as H3K9me2 (Fernandes et al. 2024). So far, the only other examples where the CO frequency increased in these pericentromeric low recombining regions was observed after performing the K.O. of the *CMT3* gene in Arabidopsis, which reduced both CHG DNA methylation and H3K9me2 (Underwood et al. 2018, Fernandes et al. 2024). However, this deregulation was at a much smaller magnitude compared to our allotriploid model. In addition, the use of the Crispr-Cas9 technology may be longer and more difficult as each gene is often present in several copies in most crops, as most of them are (paleo)polyploids (Facon et al. 2023). The natural method we present here and that plays with the ploidy level will be far more efficient for generating genetic diversity in these pericentromeric regions. This is crucial as pericentromeric regions are far from being deprived of expressed genes, including genes of interests (Boideau et al. 2021, Rousseau-Gueutin et al. 2020, Fernandes et al. 2024). For example, the *B.* napus cv. Darmor A01 and A02 pericentromeric regions contain 662 and 505 genes, representing 15.64 and 11.31% of the genes present on the A01 and A02 chromosomes, respectively. Additionally, some of these pericentromeric genes can be associated to genes involved in agronomic traits of interests, as 28 and 25 Resistant Gene Analogs were identified in these A01 and A02 pericentromeric regions, according to Rousseau-Gueutin et al. (2020). Recently, the particular interest of using Brassica allotriploidy to decrease the size of a QTLs present in pericentromeric regions was highlighted by a modelling study (Tourette al. 2021) and exemplified for a pericentromeric QTL conferring blackleg resistance (Boideau et al. 2021).

### A natural method enabling to switch on and off the strict meiosis regulation

In *Brassica*, allotriploidy was shown to deeply modify the recombination landscape (Pelé et al. 2017, Boideau et al. 2021). However, it remained to be determined whether this modification may be maintained for successive generations to take advantage of this meiotic deregulation or revert back to normal to prevent potential longer-term genomic instabilities (Vincenten et al., 2015). To test this hypothesis, we identified an AAC and AACC progenies from the first allotriploid hybrid. We were able to show that the maintenance of an allotriploid level at the second generation allows to keep this modified recombination pattern, including the presence of CO in the normally cold recombining pericentromeric regions. On the contrary, we demonstrated that recovering an allotetraploid individual can immediately switch off this deregulation. This descending allotetraploid individual recovered a normal meiotic behavior and a seed set similar to most cultivated allotetraploid *B. napus* varieties (Siles et al. 2021). The observation of the presence of a modified recombination pattern only in allotriploid but never in allotetraploid hybrids (whatever the generation) strongly indicates that interploidy (i.e. presence of one genome in single copy, where all its chromosomes remain as univalent during meiosis) is at the origin of this modification of the recombination control. Recently, the identification of a similar meiotic deregulation a wheat allopentaploid (2*n*=5*x*=35, *AABBD*; Yang et al. 2022) strongly comforts this hypothesis and suggests that this phenomenon may be common to many polyploid flowering plants. Given the important number of polyploid crops, and that many of them present an eroded genetic diversity, this modification of the recombination control via interploidy could deeply increase and shuffle their genetic diversity. Indeed, many of these polyploid crops can be crossed to their parental progenitor of a smaller ploidy level and give rise to a still fertile interploid hybrid, as exemplified in wheat (Vardi and Zohary, 1967), coffee (Krug and Mendes, 1940), strawberries (Yarnell et al. 1931), tomato (Rick et al. 1988) or tobacco (East et al. 1933).

Altogether, the easy recovery of an allotetraploid *B. napus* individual in the progeny of an allotriploid hybrid, associated with a stable meiosis and an improved seed set, is a proof of concept that the allotriploid pathway is of high relevance (i) to speed up and reduce the costs of some breeding programs by reducing the size of segregating populations and number of individuals to be genotyped, (ii) to reduce the size of *B. rapa* introgressed regions of interests (preventing the parallel introgression of deleterious alleles), (iii) and lastly to reduce the linkage disequilibrium observed within *B. napus*.

### Origin of the recombination modification

It is yet to be deciphered what factors may be at the origin of this intriguing modification of the recombination control in interploid species. One factor that may be involved in this phenomenon is the delay of meiosis progression, especially in prophase of such interploidy individual. Indeed, it has been observed that univalents move more slowly than paired chromosomes, potentially increasing the duration of meiosis (Carlton et al. 2006, Cortes et al. 2015) and therefore potentially increasing the formation of COs. However, this cannot be the only factor as there is not a linear increase of the homologous recombination rate with the number of C chromosomes in addition (Suay et al. 2014). Indeed, it has been observed that the increased homologous recombination frequency observed in such individual was mainly explained by the addition of C09, and also to a lesser extent by the C06 chromosomes (Suay et al. 2014), indicating a genetic control of this phenomenon. This observation might be related to the presence on such chromosomes of some dosage sensitive meiotic genes, as demonstrated for *HEI10* (Ziolkowski et al. 2017) and *ASY1* (Lambing et al. 2020) in *A. thaliana*, *ASY3* in *B. napus* (Chu et al. 2024) or *FIGL1* in *Z. mays* (Zhang et al. 2023), associated with a limited number of homologous chromosomes that can pair. A third factor that may be participating in this phenomenon relates to epigenetic modifications. In the monkeyflower *Mimulus*, it has notably been showed that allotriploidy was at the origin of DNA demethylation and that a partial remethylation occurred when recovering an allohexaploid (Edger et al. 2017). As observed from the K.O. of some epigenetic genes in Arabidopsis, the decrease of DNA methylation but also of some repressive histone marks, such as H3K9me2, may modify COs distribution (Melamed-Bessudo and Levy 2012, Yelina et al. 2012, Underwood et al. 2018, Fernandes et al. 2024). The potential link that may exist between DNA methylation or chromatin compaction remains to be investigated in meiotic cells of such allotriploid hybrids.

The comparison of the first- and second-generation hybrids (from the same ploidy level) revealed here the presence of a slight increase of CO frequency in the second-generation, which may be associated to the juxtaposition effect identified in Arabidopsis (Ziolkowski et al. 2015). In this latter species, it has been observed a slight increase of 1.35-fold in the heterozygous region that was juxtaposed to a homozygous region (Ziolkowski et al. 2015). The juxtaposition effect corresponds to a MSH2-dependant local redistribution of Class I COs towards polymorphic heterozygous regions to the detriment of the juxtaposed homozygous regions, in *A. thaliana* (Blackwell et al. 2020). Similarly, a slight increase of recombination in BC2 compared to BC1 hybrids was observed in *B. oleracea* (1.66-fold: Kearsey et al. 1996). Therefore, this effect may be general at least to Brassicaceae and be considered in some breeding programs.

### Putative role of an allotriploid step in the speciation success of a newly formed polyploid

Allopolyploid species may arise by numerous routes. They may form via the merging of two divergent diploid genomes through the formation of an allodiploid (i.e. homoploid bridge), allotriploid (i.e. unilateral pathway) or allotetraploid (i.e. bilateral pathway) hybrids (Tayale and Parisod, 2013, Pelé et al. 2018). In several polyploid species, only a few allopolyploidisation events were at their origin. To increase its population size, a new polyploid may produce progenies through self-fertilization or via crossing with one of its parental species. That latter route will form allotriploid individuals presenting this deregulated recombination pattern. Thereafter, the selfing of this allotriploid and/or its backcrossing with an allotetraploid individual will thus give rise to more genetically diversified allotetraploid individuals. A recent study from Cao et al. (2023) revealed that within three generations of self-fertilization, allotriploids mainly developed near complete allotetraploid individuals via gradually increasing the chromosome number and fertility. Indeed, natural selection strongly acts and favors the genomically stable allotetraploid progenies over interploid or aneuploid individuals. Nevertheless, during these few intermediate generations of aneuploidy, it is likely that these individuals also presented a modified recombination pattern (but at a lesser extent), as previously observed in *Brassica* (Suay et al. 2014). Thus, allotriploidy, with aneuploidy as intermediate, can play a major role in polyploid genetic diversification and potential adaptation, facilitating their potential establishment and speciation success. These results further support the importance of allotriploidy as a bridge in polyploid speciation.

## Material and Methods

### Plant material

The plant material analyzed in this study is described in Figure 1. To distinguish the different genomes, we used as nomenclature A_r_ and A_n_ for the A genome of *B. rapa* and *B. napus*, respectively, whereas we used C_o_ and C_n_ for the C genome of *B. oleracea* and *B. napus*, respectively. The term “Rec” as index indicates that recombination occurred on the genome concerned. To produce an allotriploid AAC and allotetraploid AACC hybrids presenting a genetically identical A genome sequence enabling an unbiased comparison of their recombination profile, we used the following strategy. To produce the allotetraploid hybrid (4x-F1), we firstly created a resynthesized oilseed rape, which was obtained by crossing the pure inbred line *B. rapa* cv. Chiifu (A_r_A_r_, 2*n*=2*x*=20) with the doubled haploid *B. oleracea* cv. HDEM (C_o_C_o_, 2*n*=2*x*=18; used as male) and then by performing embryo rescue on the obtained amphihaploid (A_r_C_o_, 2*n*=19) as described earlier (Jahier et al. 1992). This hybrid spontaneously doubled its genomes and gave rise to the resynthesized allotetraploid A_r_A_r_C_o_C_o_ (2*n*=4*x*=38), hereafter referred as ChEM. This latter plant was crossed with *B. napus* cv. Darmor (A_n_A_n_C_n_C_n_) as female, and the A_n_A_r_C_n_C_o_ F1 hybrid referred as 4x-F1 gave rise to 172 B1F1 plants after backcrossing with recurrent parent *B. napus* cv. Darmor as male (Boideau et al. 2021). One of these B1F1 plants with ~50% of A and C genome at the heterozygous stage was kept (A_Rec_A_n_C_Rec_C_n_, 2*n*=4*x*=38, referred as 4x-B1F1^4x^) to produce a B2F1 segregating population (172 plants) by backcrossing it with *B. napus* cv. Darmor as male (Figure 1).

For the allotriploid pathway, an allotriploid F1 hybrid (A_n_A_r_C_n_, 2*n*=3*x*=29, referred as 3x-F1) was obtained by crossing the allotetraploid *B. napus* cv. Darmor A_n_A_n_C_n_C_n_ (2*n*=4*x*=38) and *B. rapa* cv. Chiifu (A_r_A_r_, 2*n*=2*x*=20) as male. For the production of its B1F1 progeny, the hybrid was crossed with *B. napus* cv. Darmor as male. We selected 131 plants to generate the mapping population of the 3x-F1 hybrid (Boideau et al. 2021). Within this latter B1F1 population, we selected two AAC allotriploid hybrid progenies (A_Rec_A_n_C_n_, 2*n*=3*x*=29, hereafter referred to as 3x-B1F1a^3x^ and 3x-B1F1b^3x^) and one allotetraploid hybrid progenies (A_Rec_A_n_C_n_C_n_, 2*n*=4*x*=38, referred as 4x-B1F1^3x^) based on their meiotic behavior and on the common polymorphic regions. Then, these three hybrids were backcrossed to *B. napus* cv. Darmor as male, giving rise to three mapping populations of 130, 130 and 172 plants, respectively (Figure 1). All parental accessions were provided by the Biological Resource Center BrACySol (INRAe, Ploudaniel, France).

### DNA extraction and genotyping

Genomic DNA was extracted from lyophilized young leaves with the sbeadex maxi plant kit (LGC Genomics, Teddington Middlesex, UK) on the oKtopure robot at the GENTYANE platform (INRAe, Clermont-Ferrand, France). Genotyping data were obtained using the *Brassica* 15K and 19K Illumina Infinium SNP array (SGS TraitGenetics GmbH, Gatersleben, Germany).

To assess recombination within the pericentromeric region of A01 and A02 chromosomes that contained a very low number of polymorphic SNPs between our parental lines, we then developed an additional set of markers to specifically densify these pericentromeric regions. The improved assembly of pericentromeric regions between *B. napus* cv. Darmor v10 (Rousseau-Gueutin et al. 2020) and v5 genome assemblies (Chalhoub et al. 2014) can be visualized in Figure S1, which was obtained by comparing their assemblies using SyRI (Goel et al. 2019) with default parameters.

To design A01 and A02 pericentromeric markers, we firstly took advantage of the presence of polymorphic markers between our parental lines using data from the Brassica 60K Illumina Infinium SNP array (Clarke et al. 2016). This enabled us to identify 48 polymorph markers that were absent from the 15K and 19K Brassica Illumina Infinium arrays and for which we developed KASPar markers using their context sequences. We also designed 39 additional KASPar markers closer to the A01 and A02 centromeres by taking advantage of the whole genome assembly of the parental lines (Zhang et al. 2018, Rousseau-Gueutin et al. 2020). To design these new SNP within our intervals of interest, we firstly retrieved the genes present in single copy in each *B. napus* cv. Darmor-bzh v10 subgenome (Rousseau-Gueutin et al. 2020) and in *B. rapa* cv. Chiifu v3 (Zhang et al. 2018), preventing the putative design of markers amplifying on different paralogous regions. The orthologous copy of these genes in *B. oleracea* cv. HDEM were also retrieved using BlastP. For each of these genes, we performed an alignment of the different copies (*B. rapa*, *B. napus* copy A and copy C, *B. oleracea*) and used an in-house python script to specifically detect polymorphism between the A chromosomes of *B. napus* cv. Darmor and *B. rapa* cv. Chiifu. Overall, genotyping data were obtained for this novel set of 87 SNP markers (specific to the A01 and A02 pericentromeric regions, details for these markers are given in Table S1) and revealed by Biomark^TM^ HD system (Fluidigm technology) and KASPar^TM^ chemistry at the GENTYANE platform (INRAE, Clermont-Ferrand, France). The raw genotyping data deriving from the Brassica Illumina infinium arrays or from the KASPar technology were analyzed using either the GenomeStudio v.2011.1 (Illumina Inc., San Diego, CA, USA) or the Fluidigm SNP Genotyping Analysis v4.1.2 softwares (Wang et al. 2009). In both cases, the raw genotyping data were processed with the auto-clustering option and validated manually. The polymorphic SNPs between the parental genotypes were selected for the establishment of genetic maps. SNPs showing more than 20% of missing data and plants showing more than 25% of missing data were removed for the downstream analyses. Potential double crossovers supported by only one genetic marker and with a physical distance between these two events below 500 kb was corrected as missing data, as described in Rowan et al. (2019). To prevent the overestimation of recombination in the low recombining pericentromeric regions, plants showing more than one CO in the pericentromeric region of A01 and A02 were removed. Finally, to determine the physical position of the different markers, the context sequences of each SNP marker were physically localized on the reference genome *B. napus* cv. Darmor-bzh v10 (Rousseau-Gueutin et al. 2020) by using BlastN (ver. 2.9.0, min. e-value 1 × 10 −20, Altschul et al. 1990) and by keeping the best blast hit obtained for a given subgenome (minimum percentage of alignment and identity of 80%).

### Genetic maps

The first genetic maps were established separately for each population using the CarthaGene software (v. 1.2.3, De Givry et al. 2005). Establishment of linkage groups and SNP ordering were determined using a logarithm of odds score (LOD) threshold of 4.0 and a threshold recombination frequency of 0.3, as previously described (Pelé et al. 2017). After these few corrections, the final genetic maps were created using the Kosambi function to evaluate the genetic distances in centimorgans (cM) between linked SNP markers (Kosambi, 1943). Genotyping data used are available in Table S1. The genetic landscapes were illustrated using Circos v0.69-9 (Krzywinski et al. 2009).

The pericentromeric and centromeric borders were retrieved from Boideau et al. (2022) and were based on gene density and centromeric-specific repeats, respectively. Precisely, pericentromeric regions were defined as regions surrounding the centromere and characterized by a gene density below the chromosome average. Hereafter, a pericentromeric interval refers to the interval between the pericentromeric borders, corresponding to both the centromeric and pericentromeric regions.

### Interference

The interference parameters using the Gamma model were determined using the software CODA with the default parameters (Gauthier et al. 2011). Confidence intervals and statistical tests were not based on Fisher’s gaussian approximation as proposed in the graphical user interface of CODA but were performed based on 1000 resimulations using the command-line interface *via* custom perl and R scripts.

### Flow cytometry and cytogenetic studies

Chromosome numbers of the allotriploid progenies were assessed in leaves by flow cytometry as described in Leflon et al. (2006). Pollen viability was assessed from three independent flowers, using acetocarmine staining, as described in Jahier et al. (1992). For the establishment of meiotic behavior, samples of young floral buds were fixed in Carnoy’s II solution (alcohol:chloroform:acetic acid, 6:3:1) for 24 h at room temperature and stored until use in 50% ethanol at 4 °C. Anthers were then squashed and stained with a drop of 1% acetocarmine solution. Chromosome pairing was assessed per plant from 20 Pollen Mother Cells (PMCs) at metaphase I.

#### a) Meiotic proteins immunolocalization

The immunolabelling of HEI10 was carried out on meiotic chromosome spread prepared as described in Chelysheva et al. (2013) with minor modifications. Briefly, the chromosome preparations were incubated in Rnase A (100ng/µL) and pepsin (0.05%) in 10 mmol HCL, dehydrated in an ethanol series (70%, 90% and 100%) and air-dried. The anti-HEI10 antibody was used at a dilution of 1/100^e^ in 1X PBS-T-BSA. The slides were incubated overnight at 4°C. After three rinses in 1X PBS-T, slides were incubated for 1h at room temperature with a labeled secondary antibody (labeled goat anti-rabbit IgG Alexa fluor 488-Invitrogen (ref. A-11008, Invitrogen) diluted 1/200 in 1X PBS-T-BSA. After three rinses in 1X PBS-T, the slides were mounted in Vectashield (Vector Laboratories) containing 2.5µg/mL of 4’,6-diamidino-2-phenylindole (DAPI). Fluorescence images were captured using an ORCA-Flash4 (Hamamatsu, Japan) on an Axioplan 2 microscope (Zeiss, Oberkochen, Germany) and analyzed using Zen PRO software (version 2, Carl Zeiss, Germany).

#### b) GISH-like at meiosis using a genome specific BAC clone

For observation of HEI10 foci only on A paired chromosomes, the BAC clone BoB014O06 from *B. oleracea* (Howell et al. 2002) was used as a probe for the C genome on the same cells. This GISH-like BAC hybridized specifically to regions on every C-genome chromosome in *B. napus* (Książczyk et al, 2011). The BoB014O06 probe was labelled by random priming with biotin-14-dUTP (Invitrogen, Life Technologies). The hybridization mixture, consisting of 50% deionized formamide, 10% dextran sulfate, 2X SSC, 1% SDS and labelled probes (200ng per slide), was denatured at 92°C for 6 min and transferred to ice. The denatured probe was placed on the chromosome preparation and *in situ* hybridization was carried out overnight in a moist chamber at 37°C. After hybridization, slides were washed for 5 min in 50% formamide in 2X SSC at 42°C, followed by two washes in 4X SSC-Tween. Biotinylated probe was immunodetected by Texas Red avidin DCS (Vector Laboratories) and the signal was amplified with biotinylated anti-avidin D (Vector Laboratories). The chromosomes were mounted and counterstained in Vectashield (Vector Laboratories) containing 2.5µg/mL 4’,6-diamidino-2-phenylindole (DAPI). Fluorescence images were captured using an ORCA-Flash4 (Hamamatsu, Japan) on an Axioplan 2 microscope (Zeiss, Oberkochen, Germany) and analyzed using Zen PRO software (version 2, Carl Zeiss, Germany).

### Statistical analyses

Comparison of the crossover rates between progenies was assessed for every interval between consecutive SNP markers using a 2-by-2 chi-squared analysis considering a significance threshold of 5%. Additionally, these comparisons were also performed for a given heterozygous region, at the chromosome and at the genome scales using 2-by-2 chi-squared tests. For these tests, a conservative Bonferroni-corrected threshold of 5% was applied, using either the number of regions, the number of chromosomes or the number of intervals between adjacent SNP markers per A chromosome, or for the whole A subgenome (as described in Pelé et al. 2017 and Boideau et al. 2021). Statistical comparisons of the interference strength were assessed using the software CODA (Gauthier et al. 2011). Statistical differences for the number of HEI10 foci between the different hybrids were assessed using the Mann Whitney Wilcoxon statistical test and considering a significance threshold of 5%, with a conservative Bonferroni-corrected threshold of 5%. All statistical analyses were performed using RStudio (version 3.6.1, R Core Team, 2021).

## Acknowledgments

We thank the Genetic Resource Center BrACySol (INRAe, Ploudaniel, France, https://www6.rennes.inrae.fr/igepp_eng/About-IGEPP/Platforms/BrACySol) for providing seeds of the parental lines. We would like to thank all the staff who took care of our plant material (especially L. Charlon, P. Rolland, J-P. Constantin, J. M. Lucas and F. Letertre). We also thank the UMR 1905 GENTYANE Platform (especially C. Poncet, INRAe, Clermont-Ferrand, France, http://gentyane.clermont.inra.fr) for performing DNA extractions and KASPar genotyping. We thank SGS TraitGenetics (especially M. Ganal) for generating the genotyping data from the 15K and 19K Brassica Illumina infinium array (Gatersleben, Germany, http://www.traitgenetics.com/). We warmly thank the plant molecular cytogenetic platform (INRAe, Biogenouest, Le Rheu, France, https://www6.rennes.inrae.fr/igepp_eng/About-IGEPP/Platforms/Molecular-Cytogenetics-Platform-PCMV) for helping and participating in the cytogenetic experiments of this study. We thank Mathilde Grelon for providing the HEI10 antibodies, sharing protocols and productive discussions. We thank Eric Jenczewski for his pertinent remarks to improve the project. We are thankful to Davy Soleilhet for important insight into image analysis with the Zeiss Zen software. We thank the GOGEPP team and GenOuest bioinformatic platform (Rennes, France, https://www.genouest.org/), especially F. Legeai, S. Robin, M. Boudet and A. Bretaudeau.

## Contributions

FB, A-MC and MR-G designed the study. FB performed the genetic maps and statistical analyses. LM contributed to the automatization of the genetic map establishment. AB and VH performed the HEI10 experiments. AB, VH, GT, OC and FB analyzed the HEI10 images. GT and VH performed and analyzed the MI BAC FISH meiotic behavior. ML-T took care of the DNA samples. ML-T, GD, CF and FB analyzed the genotyping data. FE performed the classical meiotic behavior. MG generated and took care of the plant material. JB designed the additional pericentromeric SNP. JM generated the Circos representations. MF and OM analyzed the interference. FB, MR-G and AM-C wrote the manuscript. All authors approved the final version of the manuscript.

## Data availability

All data analyzed in this study are provided as supplementary material.

## List of Supplementary Tables

Table S1: Sequences of the KASPar markers used

Table S2: Genotyping data used to generate the generic maps Table S3: HEI10 foci quantification used for Figure 4

## List of Supplementary Figures

Figure S1: Improvement of the genome assembly of *B. napus* cv. Darmor

