## Supplementary figures and images for "Playing with the ploidy level enables to switch on and off the strict recombination control even in the vicinity of *Brassica* centromeres"

### Figure S1

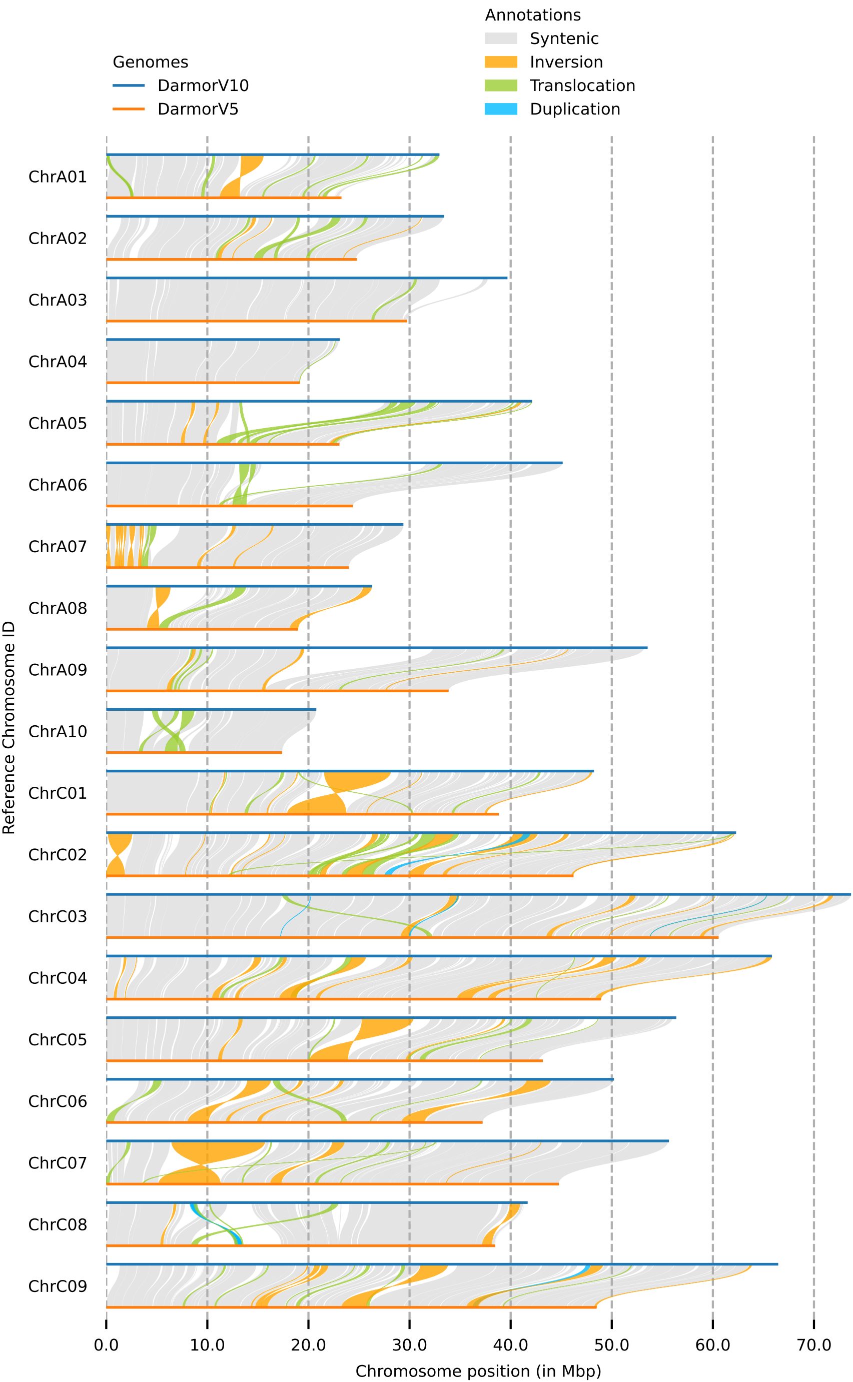
